# Confinement prevents postmenopausal osteoporosis and obesity via nuclear mechanical remodeling

**DOI:** 10.64898/2025.12.15.693823

**Authors:** Min Xu, Lei Wang, Weihao Wang, Haoxiang Yang, Hongyuan Jiang

## Abstract

Postmenopausal osteoporosis (PMOP), affecting about half of postmenopausal women, is a major global health burden characterized by fragile bones and frequent comorbidity with obesity. Bone marrow mesenchymal stem cells (BMSCs) differentiation imbalance underpins PMOP pathogenesis and represents an attractive therapeutic target. Here, we demonstrate mechanically confining BMSCs to soft substrates mimicking physiological bone marrow stiffness completely suppresses adipogenic differentiation while inducing osteogenic differentiation. Mechanistically, confinement-induced elevation of nuclear envelope tension upregulates Lamin A/C expression, which in turn modulates Runx2 nuclear localization by activating histone acetylation via KAT2B, thereby directing BMSCs osteogenesis. Remarkably, FDA-approved deacetylation inhibitor Chidamide significantly enhances bone regeneration while concurrently suppressing obesity in a PMOP animal model, highlighting the significant therapeutic potential of nuclear mechanotransduction as a target for PMOP and obesity treatment. This work not only provides new insights into the mechanical regulation of bone homeostasis but also opens new avenues for the treatment of osteoporosis-related disorders.

## Introduction

Postmenopausal osteoporosis (PMOP) as a globally prevalent skeletal disorder, is characterized by compromised bone strength in affected individuals, with a prevalence of fragility fractures as high as ∼40% [1, 2]. The PMOP is often accompanied by the development of obesity [3, 4], severely impacting patients’ quality of life [5]. Exacerbated by the aging population, approximately 50% of postmenopausal women suffer from PMOP [5, 6], thereby posing a serious challenge to the public health. The pathogenesis of PMOP stems from dysregulated bone remodeling, specifically characterized by reduced osteoblast differentiation and pathological expansion of bone marrow adipose tissue [5]. In essence, this reflects an imbalance in the osteogenic and adipogenic differentiation of bone marrow mesenchymal stem cells (BMSCs) [7–9].

Current pharmacotherapies for PMOP primarily target osteoclast differentiated from hematopoietic stem cells, offering transient suppression of bone resorption but exhibiting high relapse rates upon treatment cessation [10]. To address this limitation, emerging therapeutic strategies focus on rebalancing the osteogenic-adipogenic differentiation of BMSCs to achieve sustained disease management. For example, teriparatide, a pharmacological agent targeting parathyroid hormone (PTH), counteract PMOP by stimulating osteoblast differentiation to enhance bone formation [11, 12]. However, its clinical utility is constrained by suboptimal efficacy, mandatory subcutaneous administration, and potential adverse effects, including aberrant bone vascularization and possible elevated oncogenic risks [1, 13]. These challenges underscore the urgent need to elucidate the *in vivo* regulatory mechanisms governing BMSCs lineage commitment and to develop safer, more effective treatment modalities.

Emerging evidence indicates that BMSC differentiation is governed not only by biochemical signals, such as hormones and cytokines, but also by mechanical cues derived from the extracellular matrix (ECM) [14]. For instance, a well-known experimental observation is that rigid substrates (22–100 kPa) promote osteogenic differentiation, whereas soft substrates (0.5-3 kPa) promote adipogenic differentiation [15–17]. Although substrate stiffness has been widely recognized as a key regulator of mesenchymal stem cells fate *in vitro* [15, 16], this paradigm fails to fully account for BMSCs differentiation behavior *in vivo*. Specifically, although physiological bone marrow tissue exhibits a soft mechanical signature with stiffness ranging from 10^-1^ to 10^0^ kPa [18], it still supports robust osteogenic differentiation, suggesting that additional biomechanical or contextual factors may contribute to lineage commitment beyond stiffness.

Notably, osteogenically differentiated BMSCs exhibit a distinct spatial preference, predominantly localizing to perivascular niches adjacent to the endosteum [19, 20]. These regions, situated in the cortical bone are characterized by high cellular density, subjecting BMSCs undergoing osteogenic differentiation to substantial compressive forces from neighboring cells and the surrounding ECM, effectively creating confined microenvironments.

Mounting evidence indicates that crowding-induced confinement profoundly impacts cell behaviors and functions [21, 22]. For example, cellular crowding can lead to the accumulation of mechanical stress, thereby reducing cellular proliferation [21], causing cell extrusion [23], and inhibiting multipolar division [24]. Confinement can also drive cancer cell migration via directed water and ion transport in very narrow channels [25], and induce the mesenchymal-amoeboid transition under the condition of low adhesion [26–28]. Intriguingly, confinement can even make initially quiescent round cells on soft adhesive substrates spread and migrate, similar to the phenotype on unconfined stiff substrates [29]. Based on these observations, we propose that spatial confinement represents a previously underappreciated yet critical biomechanical cue governing BMSCs lineage specification *in vivo*, and it may contribute substantially to the osteogenic-adipogenic imbalance underlying PMOP, offering new insights into its pathogenesis and potential therapeutic strategies.

Here, we found that the crowding microenvironments in human femoral tissue is highly correlated with pathological features of PMOP, and further demonstrate that spatial confinement completely abolishes adipogenic differentiation and induces osteogenic differentiation in BMSCs cultured on soft substrates. We revealed that confinement promotes nuclear deformation, increases nuclear envelope tension, and consequently drives the remodeling of Lamin A/C. This mechanical remodeling of the nuclear lamina modulates the nuclear localization of the osteogenic transcription factor Runx2 by activating deacetylase via KAT2B (Lysine Acetyltransferase 2B), thereby directing BMSCs osteogenesis. Leveraging this mechanotransduction pathway, we employed Chidamide—an orally available deacetylase inhibitor approved by FDA—to feed the ovariectomy (OVX) model rats. Strikingly, Chidamide treatment significantly enhanced bone regeneration while concurrently suppressing obesity, a hallmark comorbidity of PMOP. Our findings identify confinement as a critical mechanical regulator of BMSCs lineage specification *in vivo,* and identifies mechanical confinement-induced epigenetic remodeling as a promising target for PMOP. This discovery not only advances our understanding of bone homeostasis, but also opens new avenues for treating osteoporosis and related disorders.

## Result

### Crowding-induced spatial confinement, which fully overrides the well-established effects of both ECM stiffness and biochemical cues, can directly drive the adipogenic-to-osteogenic differentiation transition

To investigate the correlation between crowding and differentiation of BMSCs within the bone marrow cavity *in vivo*, we harvested femur and bone marrow tissue slices from osteoporosis patients and healthy controls (Fig. 1a) for H&E staining (Fig. 1b-d) and mechanical characterization (Fig. 1e). We found that adipocyte diameter (Fig. 1c) and volume fraction (Fig. 1d) significantly increased in the femur tissue of osteoporosis patients, consistent with previous studies [30, 31] that osteoporosis is often accompanied by an increase in bone marrow adipose tissue. Notably, we observed that both osteoporotic and healthy bone marrows are very soft, and there is no significant difference between their stiffness (Fig. 1e), indicating the bone marrow stiffness is not the factor affecting the imbalance in the osteogenic and adipogenic differentiation of BMSCs. In contrast, the healthy control group exhibited markedly higher cell density and less adipocyte compared to osteoporotic group (Fig. 1f-g), suggesting there is some correlation between crowding and osteogenic differentiation.

**Fig. 1.**
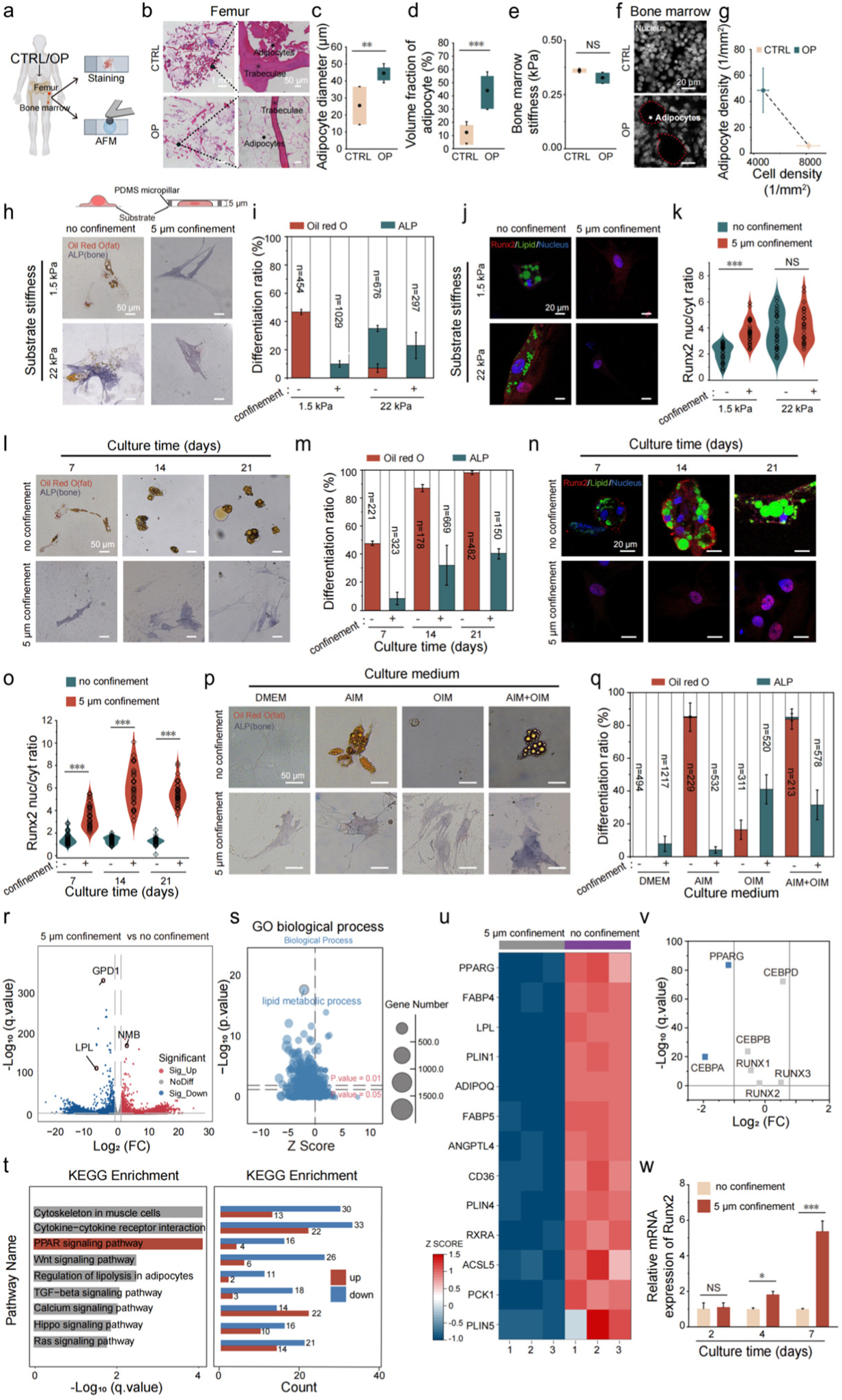
Crowding-induced spatial confinement, which fully overrides the well-established effects of both ECM stiffness and biochemical cues, can directly drive the adipogenic-to-osteogenic differentiation transition. **(A)** Schematic showing the stiffness measurement and the staining analysis of bone tissues in healthy controls or osteoporosis donors. **(B-D)** HE staining (B) and quantitative analysis (C-D) of adipocytes of femur in healthy controls or osteoporosis donors (each n=3, 2 biologically independent experiments in each sample). **(E)** The bone marrow stiffness of healthy controls or osteoporosis donors (each n=3, 10 positions were randomly measured to take the mean in each sample). **(F-G)** The nuclear staining images of bone marrow biopsy (F) and the correlation between cell density and adipocytes (G, each n=3, 2 biologically independent experiments in each sample). **(H-I)** Bright-field images (H) and quantitative analysis (I) of BMSCs with or without 5 μm confinement, cultured on soft/stiff substrates (1.5/22 kPa) differentiated in osteogenic-adipogenic mixed medium: oil red O (fat) and ALP (osteogenic) double staining for 3 biologically independent experiments. **(J-K)** Immunofluorescence staining (J) and quantification (K, each n=31 cells) of Runx2 (red, osteogenic marker), LipidTOX^TM^ (green, lipid droplets) and nucleus (blue) of BMSCs induced differentiation on soft/stiff substrates. **(L-M)** Bright-field staining (L) and quantitative analysis (M, 3 biologically independent experiments in each group) of BMSCs’ differentiation with different culture time in mixed induction medium. (N-O) Immunofluorescence staining (N) and quantitative analysis (O, each n=31 cells) of Runx2 (red, osteogenic marker), LipidTOX^TM^ (green, lipid droplets) and nucleus (blue) of BMSCs with different culture time in mixed induction medium. **(P-Q)** Bright-field staining (P) and comparison (Q, 3 biologically independent experiments in each group) of BMSCs differentiation efficiency under different culture conditions. DMEM: Dulbecco’s Modified Eagle Medium; AIM: Adipogenesis induction medium; OIM: Osteogenic induction medium; AIM+OIM: Adipogenesis induction medium osteogenic induction medium volume ratio1:1. (R) Volcano map (R) of differentially expressed genes (|log2(FC)|>1, -log10(q.value)>0). **(S-T)** GO biological process (S) KEGG pathway enrichment analysis (T) of BMSCs with 5 μm confinement (p<0.05). **(U-V)** Heat map (U) of differentiation-related genes and volcano gram (V) of osteogenic/adipogenic core gene expression of BMSCs with or without 5 μm confinement (each n=3). (W) Runx2 mRNA temporal expression of BMSCs with 5 μm confinement on soft substrates (3 independent qPCR assays). Data: C, D, E, I, K, M, O, Q, W, mean ± SD; G mean ± SE; Tukey’s test: ns, P > 0.05; *P ≤ 0.05; **P ≤ 0.01; ***P ≤ 0.001.

To mimic the mechanical confinement induced by crowding *in vitro*, we constrain the cell height of BMSCs (Fig. 1h and Supplementary Fig. 1a-b) without affecting cell viability (Supplementary Fig. 1c-d) and changing the material properties including porosity (Supplementary Fig. 2a), stiffness (Supplementary Fig. 2b) and viscoelasticity (Supplementary Fig. 2c-k) of the substrate. The Oil red O and alkaline phosphatase (ALP) staining experiments (Fig. 1h) indicated that under the conditions of mixed adipogenic induction medium (AIM) and osteogenic induction medium (OIM), unconfined BMSCs on soft substrates (1.5 kPa) undergo adipogenic differentiation (Fig. 1h-i), consistent with previous studies [15–17]. Strikingly, we found that confined BMSCs undergo osteogenic differentiation without any adipogenic differentiation (Fig. 1h-i). In fact, if not confined, a small fraction of BMSCs can still undergo adipogenic differentiation even on stiff substrates with the optimal stiffness (22 kPa) for osteogenic differentiation (Fig. 1h-i). However, under confinement, adipogenic differentiation completely disappeared on both stiff and soft substrates (Fig. 1h-i). Immunofluorescence staining suggested that confinement promoted the nuclear translocation of Runx2 (a master regulator of osteogenesis) in BMSCs on soft substrates (Fig. 1j-k), further confirming that confinement can promote osteogenesis. Furthermore, the confinement-induced adipogenic-to-osteogenic differentiation transition was continuously enhanced with the extension of the culture time (Fig. 1l-o). After 21 days of culture, the adipogenic differentiation ratio of unconfined BMSCs increased from 47.5% (observed at day 7) to 98%, and no osteogenic differentiation occurred (Fig. 1m). In contrast, the adipogenic differentiation ratio of confined BMSCs remained 0% and the osteogenic differentiation ratio of confined BMSCs increased from 8.8% to 40.8% (Fig. 1m). Even in the case of adipogenic induction medium (AIM) and DMEM, confinement can still promote osteogenesis while completely abolishing adipogenesis (Fig. 1p-q), indicating this confinement-induced adipogenic-to-osteogenic differentiation transition does not rely on the culture medium and the impact of crowding-induced spatial confinement can fully override the well-established effects of both ECM stiffness and biochemical cues in this case.

To investigate the effect of confinement on the expression of differentiation-related genes, we performed RNA sequencing on BMSCs from the confinement group and control group after 48 hours culture on soft substrates in mixed culture medium (AIM+OIM). Confinement treatment significantly induces changes in intracellular expression profiles compared with unconfined cells (Fig. 1r). The Go Biological Process analysis of the differential genes showed that the main biological function of these genes was lipid metabolism process (Fig. 1s). Through KEGG enrichment (Fig. 1t) and core gene expression analysis (Fig. 1u-v), we found that the gene expression of the PPARγ signaling pathway related to adipogenic differentiation in confined BMSCs was generally downregulated, which is consistent with the confinement-induced inhibition of adipogenesis discovered above (Fig. 1h-q). Furthermore, RNA-seq analysis (Fig. 1v) and qPCR experiment (Fig. 1w) revealed no significant difference in Runx2 mRNA levels within the first 48 hours (Fig. 1n-o), suggesting that the initial osteogenic commitment is transcription-independent. However, a sustained increase in Runx2 expression was observed at later time points (Fig. 1w). Together with data demonstrating enhanced Runx2 nuclear localization (Fig. 1j-k and 1n-o), these findings indicate that mechanical confinement regulates osteogenesis through a biphasic mechanism: it initially promotes the nuclear translocation and functional activation of pre-existing Runx2, and subsequently induces a transcriptional upregulation of Runx2 to reinforce and maintain the osteogenic phenotype. Collectively, these data suggest that crowding-induced spatial confinement can drive the transition from adipogenic differentiation to osteogenic differentiation of BMSCs on soft substrates similar to bone marrow stiffness.

### Confinement-induced adipogenic-to-osteogenic differentiation transition is dependent on nuclear envelope tension rather than cell-substrate adhesion

Previous studies have suggested that cellular sensing of mechanical cues in the extracellular microenvironment is largely dependent on cell-substrate adhesion [32, 33]. To verify this possibility, we treated cells with 10 μM PF-573228 (a focal adhesion kinase inhibitor) and 10 μM Cilengitide (an integrin protein inhibitor) to inhibit cell-substrate adhesion (Supplementary Fig. 3a-c). No differences were found between the PF-573228-treated, Cilengitide-treated and control groups in either the Runx2 nucleus/cytoplasm ratio (Fig. 2a-b) or the proportion of ALP-positive cells (Fig. 2a-c), indicating that confinement-induced osteogenic differentiation does not depend on cell-substrate adhesion.

**Fig. 2:**
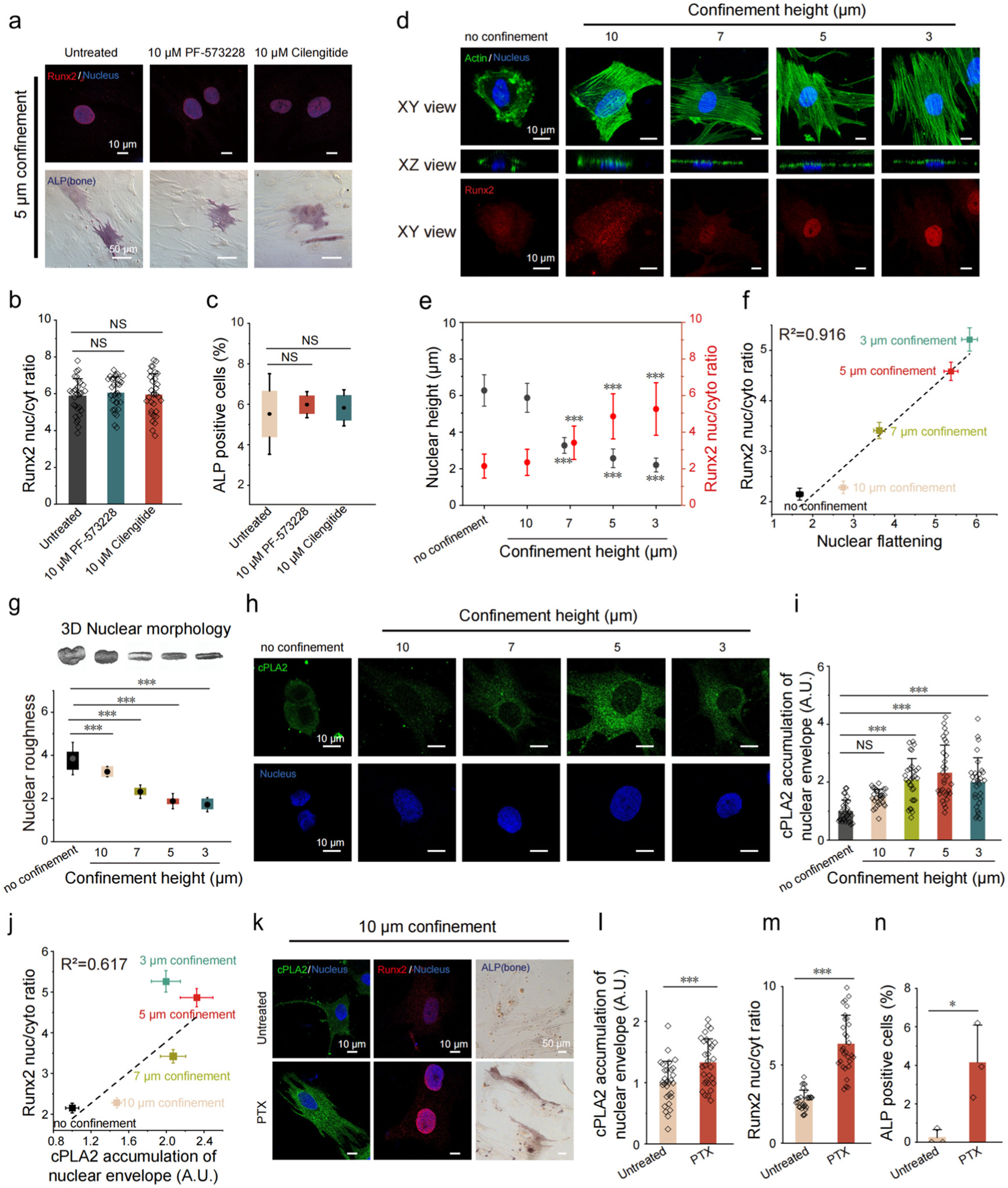
Confinement-induced adipogenic-to-osteogenic differentiation transition is dependent on nuclear envelope tension rather than cell-substrate adhesion. (**A**) Immunofluorescence staining (top panel) and ALP staining Bright-field images (bottom panel) of confined BMSCs with or without 10 μM PF-573228 (a FAK inhibitor) or 10 μM Cilengitide (an integrin inhibitor) treatment on soft substrates. (**B-C**) Runx2 nucleus/cytoplasm ratio (B, each n=31 cells) and percentage of ALP-positive cells (C, 3 biologically independent experiments in each group) of confined BMSCs with or without 10 μM PF-573228 (a FAK inhibitor) or 10 μM Cilengitide (an integrin inhibitor) treatment on soft substrates. (**D**) Images of F-actin (green), nucleus (blue) and Runx2 (red) of BMSCs on soft substrates under various degree of confinement. (**E**) Statistical results of nuclear height and Runx2 nucleus/cytoplasm ratio of BMSCs with various confinement heights (each n=30 cells). (**F**) Correlation between Runx2 nucleus/cytoplasm ratio and nuclear flattening index (R^2^=0.916, each n=30 cells). (**G**) Nuclear roughness and corresponding 3D nuclear morphology of BMSCs on soft substrates under various levels of confinement (each n=26 cells). (**H-I**) Immunofluorescence staining (H) and quantification (I, each n=30 cells) of cPLA2 at nuclear envelope of BMSCs on soft substrates under various degree of confinement. (**J**) Correlation between Runx2 nucleus/cytoplasm ratio and cPLA2 accumulation of nuclear envelope (R^2^=0.617, each n=30 cells). (**K-N**) Images (K) and quantification of cPLA2 (L, each n=30 cells), Runx2 (M, each n=30 cells) and ALP (N, 3 biologically independent experiments in each group) at nuclear envelope of BMSCs with or without 20 nM PTX treatment under 10 μm confinement. Data: B, C, E, G, I, L, M, N, mean ± SD; F, J, mean ± SE, Linear Fit; Tukey’s test: ns, P > 0.05; *P ≤ 0.05; **P ≤ 0.01; ***P ≤ 0.001.

Since our previous study has revealed the nucleus can also be used as an independent mechanical sensor to trigger confinement-induced cell spreading and migration [29], we speculate that the confinement-induced adipogenic-to-osteogenic differentiation transition may also depend on the nucleus. To test this hypothesis, we measured the Runx2 nuclear translocation under various levels of confinement (10, 7, 5 and 3 μm) on soft substrates (Fig. 2d). We found that the Runx2 nucleus/cytoplasm ratio increased only when the nucleus was significantly deformed (Fig. 2d-e and Supplementary Fig. 3d-e). Notably, the deformation of confined nuclei can be characterized by nuclear flattening [29, 34], which highly correlated with the Runx2 nucleus/cytoplasm ratio (Fig. 2f). Furthermore, the surface roughness of the nucleus reduced as the confinement height decreased (Fig. 2g), indicating the increase of nuclear envelope (NE) tension indued by confinement. Therefore, we infer that the confinement-induced adipogenic-to-osteogenic differentiation transition might be intrinsically induced by the NE tension.

To test this hypothesis, we quantified the phosphorylated form of the nucleoprotein cPLA2, as NE tension can mediate the translocation of this protein to NE [27, 28, 35]. The localization of cPLA2 at NE increased with the degree of confinement (Fig. 2h-i), indicating NE tension was indeed elevated after spatial confinement. Moreover, the cPLA2 accumulation of NE was also highly correlated with osteogenic differentiation (Fig. 2j). To further verify this hypothesis, we treated BMSCs with 20 nM paclitaxel (Taxol) for 72 hours, which can produce polyploid DNA, make the nucleus expand and then stretch NE, thereby increasing the NE tension [29]. After treatment of paclitaxel, the localization of cPLA2 at NE (Fig. 2k-l), the Runx2 nucleus/cytoplasm ratio (Fig. 2m), and the proportion of ALP-positive cells (Fig. 2n) significantly increased under the condition of 10 μm confinement compared to the untreated groups, confirming that the NE tension plays a critical role in triggering the Runx2 nuclear translocation and osteogenic differentiation.

### Lamin A/C expression mediated by NE tension can promote osteogenic differentiation in BMSCs

To explore the effects of NE tension in nuclear mechanical remodeling, we further analyzed the RNA sequencing results and observed a significant upregulation of Lamin A/C in the confinement group, with its expression level approximately twice that of control group (Fig. 3a), consistent with previous studies [35–38]. This transcriptional induction was also corroborated at the protein level. As the confinement height decreased, we observed a significant upregulation of nuclear Lamin A/C (Fig. 3b-c). The strong correlation between Lamin A/C expression and cPLA2 accumulation (Fig. 3d) suggests the regulatory role of NE tension. On the other hand, the highly positive correlation between Lamin A/C expression and Runx2 nuclear translocation (Fig. 3e) suggests that Lamin A/C may have its potential regulatory function in osteogenic differentiation. Furthermore, the immunofluorescence analysis revealed that the Runx2 nucleus/cytoplasm ratio increased progressively with Lamin A/C expression (Fig. 3f-g), suggesting a mechanosensitive coupling between NE tension, Lamin A/C expression, and Runx2 nuclear translocation.

**Fig. 3:**
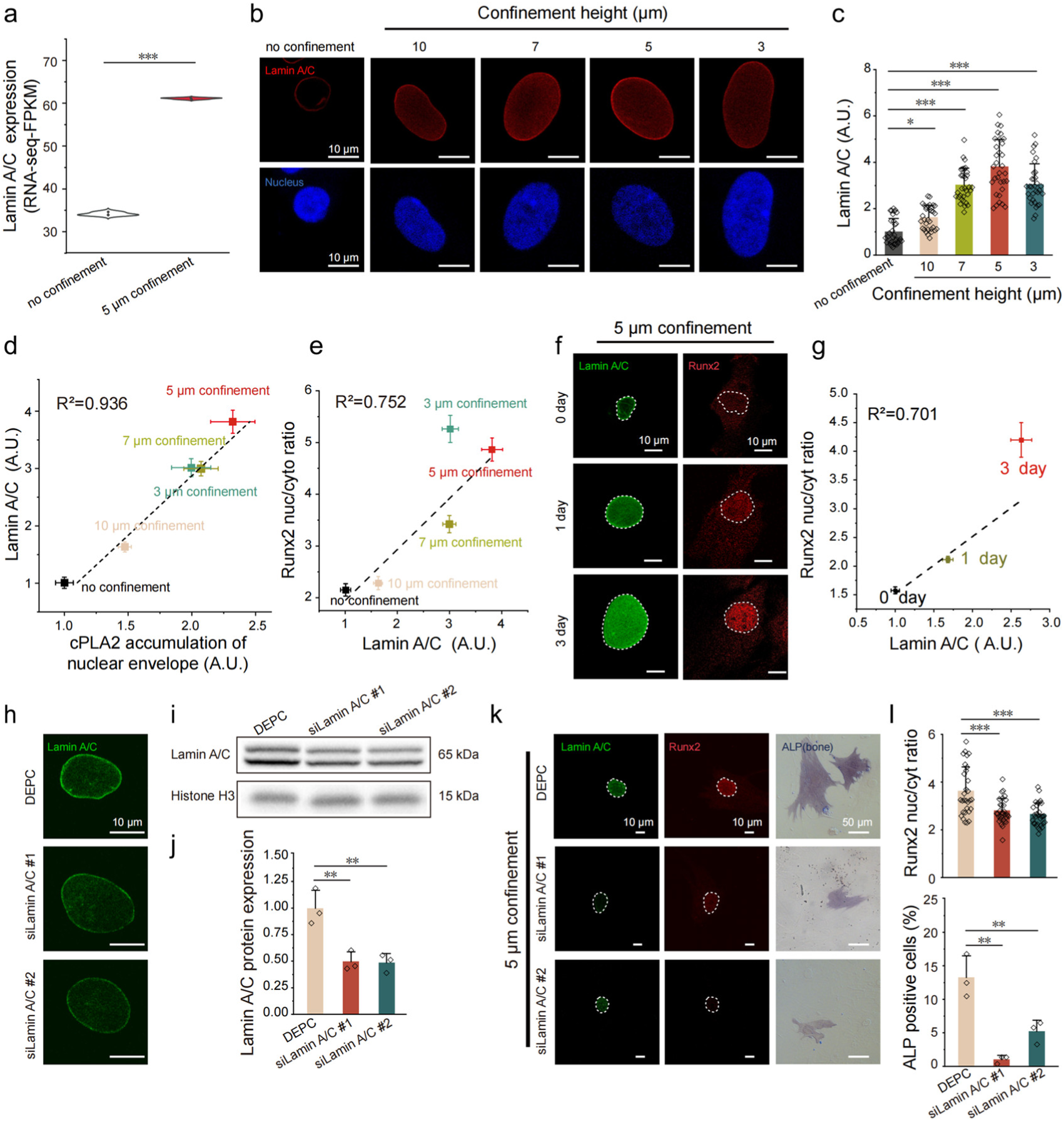
Lamin A/C expression mediated by NE tension can promote osteogenic differentiation in BMSCs. (**A**) Lamin A/C expression level of BMSCs with or without 5 μm confinement from RNA-seq analysis (FPKM, Fragments Per Kilobase Million). (**B-C**) Immunofluorescence staining (B) and quantification (C, each n=32 cells) of Lamin A/C expression of BMSCs with different confinement height on soft substrates. (**D**) Correlation between Lamin A/C expression level and cPLA2 accumulation of nuclear envelope (R²=0.936, each n=30 cells). (**E**) Correlation between Lamin A/C expression level and Runx2 nucleus/cytoplasm ratio (R²=0.752, each n=30 cells). (**F**) Images of Lamin A/C (green) and Runx2 (red) of BMSCs with 5 μm confinement on soft substrates after different culture time. (**G**) Correlation between Lamin A/C expression level and Runx2 nucleus/cytoplasm ratio (R²=0.701, each n=29 cells). (**H-J**) Immunofluorescence staining (H), Western blot (I, Histone H3 internal control) and quantification (J) of Lamin A/C knockdown verification after 3 days of siLamin A/C #1 or siLamin A/C #2 treatment (3 biologically independent experiments in each group). (**K**) Immunofluorescence images of Lamin A/C (green) and Runx2 (red) staining, and Bright-field images of BMSCs in DEPC, siLamin A/C #1 and siLamin A/C #2 group under 5 μm confinement on soft substrates. (**L**) Runx2 nucleus/cytoplasm ratio (top panel, each n=30 cells) and percentage of ALP-positive cells (bottom panel, 3 biologically independent experiments in each group) of BMSCs in DEPC, siLamin A/C #1 and siLamin A/C #2 group under 5 μm confinement on soft substrates. Data: A, C, J, L, mean ± SD; D, E, G, mean ± SE, Linear Fit; Tukey’s test: ns, P > 0.05; *P ≤ 0.05; **P ≤ 0.01; ***P ≤ 0.001.

To functionally validate Lamin A/C’s role, we performed siRNA-mediated knockdown (siLamin A/C #1 and siLamin A/C #2), confirming reduced Lamin A/C expression via Western blot and immunofluorescence (Fig. 3h-j). Notably, Lamin A/C depletion significantly attenuated Runx2 nuclear translocation and decreased the proportion of ALP-positive cells (Fig. 3k-l), indicating impaired confinement-induced osteogenic differentiation. Collectively, these findings demonstrate that confinement-induced NE tension upregulates Lamin A/C, which in turn promotes osteogenic commitment by facilitating Runx2 nuclear accumulation. This mechanotransduction pathway highlights Lamin A/C as a critical mediator of BMSCs differentiation in response to mechanical cues.

### Mechanical confinement promotes osteogenic differentiation via KAT2B-mediated histone acetylation

To elucidate the mechanism by which Lamin A/C enhances osteogenic differentiation, we systematically analyzed confinement-induced transcriptional changes in BMSCs from RNA sequencing data, identifying KAT2B (a histone acetyltransferase) as a potential downstream effector of Lamin A/C in differentiation modulation (Fig. 4a). We found that mechanical confinement significantly upregulated KAT2B expression (Fig. 4b-c), accompanied by elevated histone acetylation levels (Fig. 4b-d). Lamin A/C knockdown attenuated confinement-induced KAT2B upregulation (Fig. 4e-f) and correspondingly diminished acetylation levels (Fig. 4g-h), confirming KAT2B as a downstream effector of Lamin A/C in acetylation regulation. Genetic knockdown KAT2B with siKAT2B (Fig. 4i-n) or pharmacological inhibition of KAT2B with PCAF-IN-1 [39] (Supplementary Fig. 4a-d) both significantly attenuated histone acetylation, Runx2 nucleus/cytoplasm ratio and the proportion of ALP-positive cells in confined BMSCs, indicating the impaired osteogenic commitment. Therefore, these data establish KAT2B as a mechanoresponsive epigenetic regulator downstream of Lamin A/C, essential for mediating confinement-induced osteogenic differentiation.

**Fig. 4:**
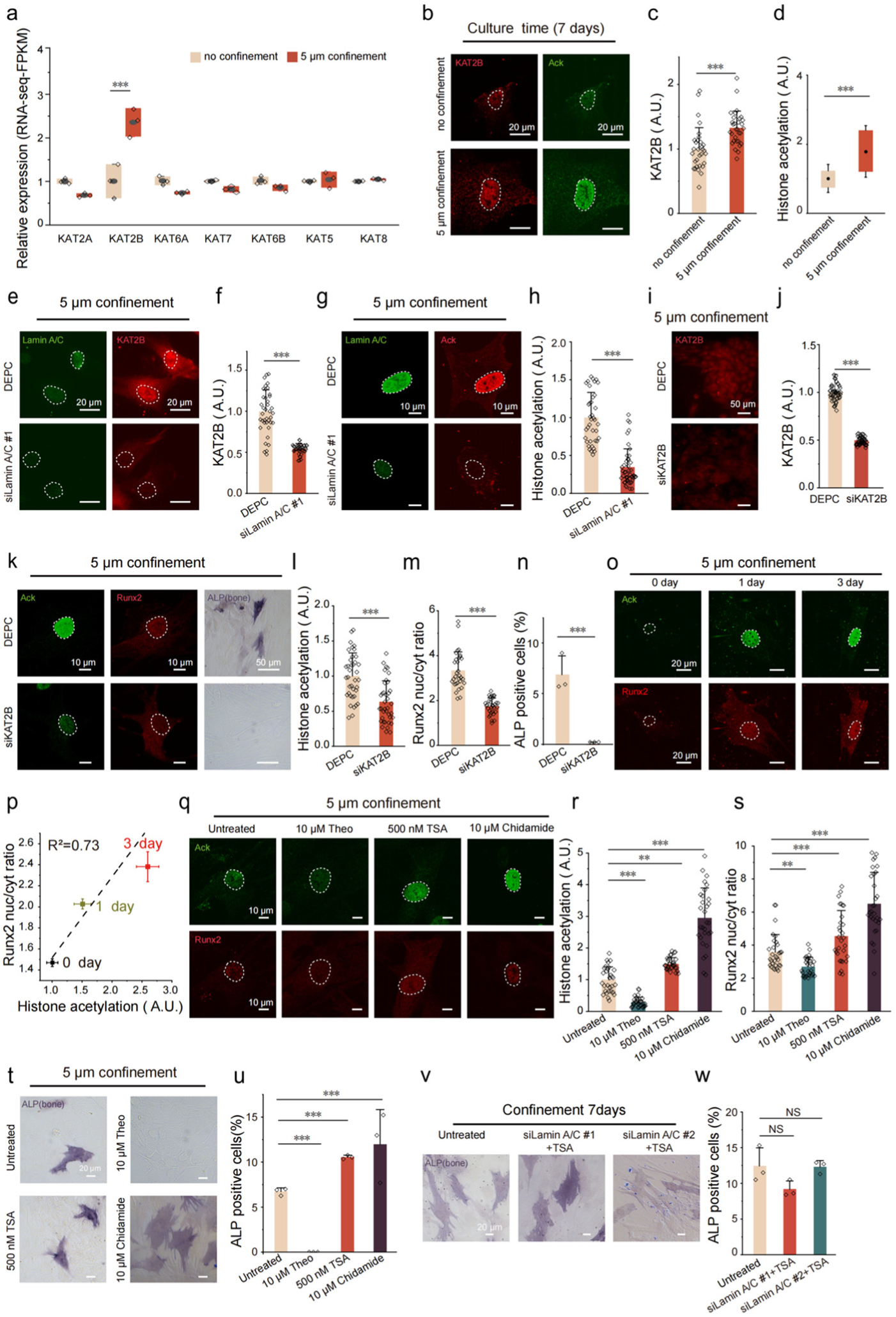
Mechanical confinement promotes osteogenic differentiation via KAT2B-mediated histone acetylation. (**A**) RNA-seq analysis of BMSCs with or without 5 μm confinement (FPKM values for acetyltransferases). (**B**) Immunofluorescence images of KAT2B (red) and Ack (green) of BMSCs with or without 5 μm confinement on soft substrates. (**C-D**) Immunofluorescence intensity quantification of KAT2B (C) and Histone acetylation (D) of BMSCs with or without 5 μm confinement on soft substrates (each n=31 cells). (**E-F**) Immunofluorescence images (E) and quantification (F, each n=39 cells) of KAT2B of BMSCs with DEPC and siLamin A/C#1 group under 5 μm confinement on soft substrates. (**G-H**) Immunofluorescence images (G) and quantification (H, each n=38 cells) of Ack of BMSCs with DEPC and siLamin A/C #1 group under 5 μm confinement on soft substrates. (**I-J**) Immunofluorescence images (I) and quantification (J, each n=47 cells) of KAT2B of BMSCs with DEPC and siLamin A/C #1 group under 5 μm confinement on soft substrates. (**K**) Immunofluorescence images of ACK (green) and Runx2 (red), and Bright-field images of ALP (osteogenic) staining of BMSCs with DEPC and siKAT2B group under 5 μm confinement on soft substrates. (**L-N**) Quantification of Histone acetylation (L, each n=38 cells), Runx2 nucleus/cytoplasm ratio (M, each n=31 cells) and percentage of ALP-positive cells (N, 3 biologically independent experiments in each group) of BMSCs with DEPC and siKAT2B group under 5 μm confinement on soft substrates (each n=38 cells). (**O**) Images of Ack (green) and Runx2 (red) of BMSCs with 5 μm confinement on soft substrates after different culture time. (**P**) Correlation between Histone acetylation expression and Runx2 nucleus/cytoplasm ratio (R²=0.73, each n=29 cells). (**Q**) Images of ACK (green) and Runx2 (red) of untreated BMSCs or BMSCs treated with 10 μM Theo (Theophylline), 500 nM TSA (Trichostatin A) and 10 μM Chidamide under 5 μm confinement on soft substrates. (**R-S**) Quantification of Histone acetylation (R) and Runx2 nucleus/cytoplasm ratio (S) of untreated BMSCs or BMSCs treated with 10 μM Theo, 500 nM TSA and 10 μM Chidamide under 5 μm confinement on soft substrates (each n=30 cells). (**T-U**) ALP (osteogenic) staining (T) and percentage of ALP-positive cells (U, 3 biologically independent experiments in each group) of untreated BMSCs or BMSCs treated with 10 μM Theo, 500 nM TSA and 10 μM Chidamide under 5 μm confinement on soft substrates. (**V-W**) ALP (osteogenic) staining (V) and percentage of ALP-positive cells (W, 3 biologically independent experiments in each group) of BMSCs with control, siLamin A/C#1+TSA and siLamin A/C#2+TSA group under 5 μm confinement on soft substrates. Data: A, C, D, F, H, J, L, M, N, R, S, U,W, mean ± SD; P, mean ± SE, Linear Fit; Tukey’s test: ns, P > 0.05; *P ≤ 0.05; **P ≤ 0.01; ***P ≤ 0.001.

Strikingly, histone acetylation levels exhibited a strong positive correlation with Runx2 nuclear accumulation in confined cells over time (Fig. 4o-p). To further dissect the connection between histone acetylation and confinement-induced osteogenic differentiation, we used 10 μM Theophylline (Theo) to inhibit histone acetylation and 500 nM Trichostatin A (TSA) or 10 μM Chidamide to enhance histone acetylation (Fig. 4q-r and Supplementary Fig. 4e-h). Inhibition of histone acetylation suppressed the Runx2 nuclear localization (Fig. 4s) and completely terminated the confinement-induced osteogenic differentiation (Fig. 4t-u), whereas enhancement of histone acetylation promoted both (Fig. 4s-u). Notably, rescue experiments further demonstrated that TSA treatment restored osteogenic capacity in siLamin A/C BMSCs under mechanical confinement (Fig. 4v–w). Collectively, these findings demonstrate that mechanical confinement promotes osteogenic differentiation through KAT2B-dependent histone acetylation.

### Pharmacological enhancement of histone acetylation effectively mitigates osteoporosis and obesity progression *in vivo* via nuclear mechanotransduction signaling

To evaluate the therapeutic potential of histone acetylation in osteogenic differentiation regulation *in vivo*, we established a bilateral ovariectomy (OVX) rat model mimicking PMOP [40–42], with cohorts assigned to Sham, OVX, and OVX + Chidamide groups. After two weeks of postoperative recovery, nine weeks of oral gavage administration were implemented (Fig. 5a). Subsequent assessment of femoral bone marrow stiffness showed no significant intergroup differences (p>0.05) (Fig. 5b), indicating that neither surgical procedures nor pharmacological treatments altered bone marrow stiffness.

**Fig. 5:**
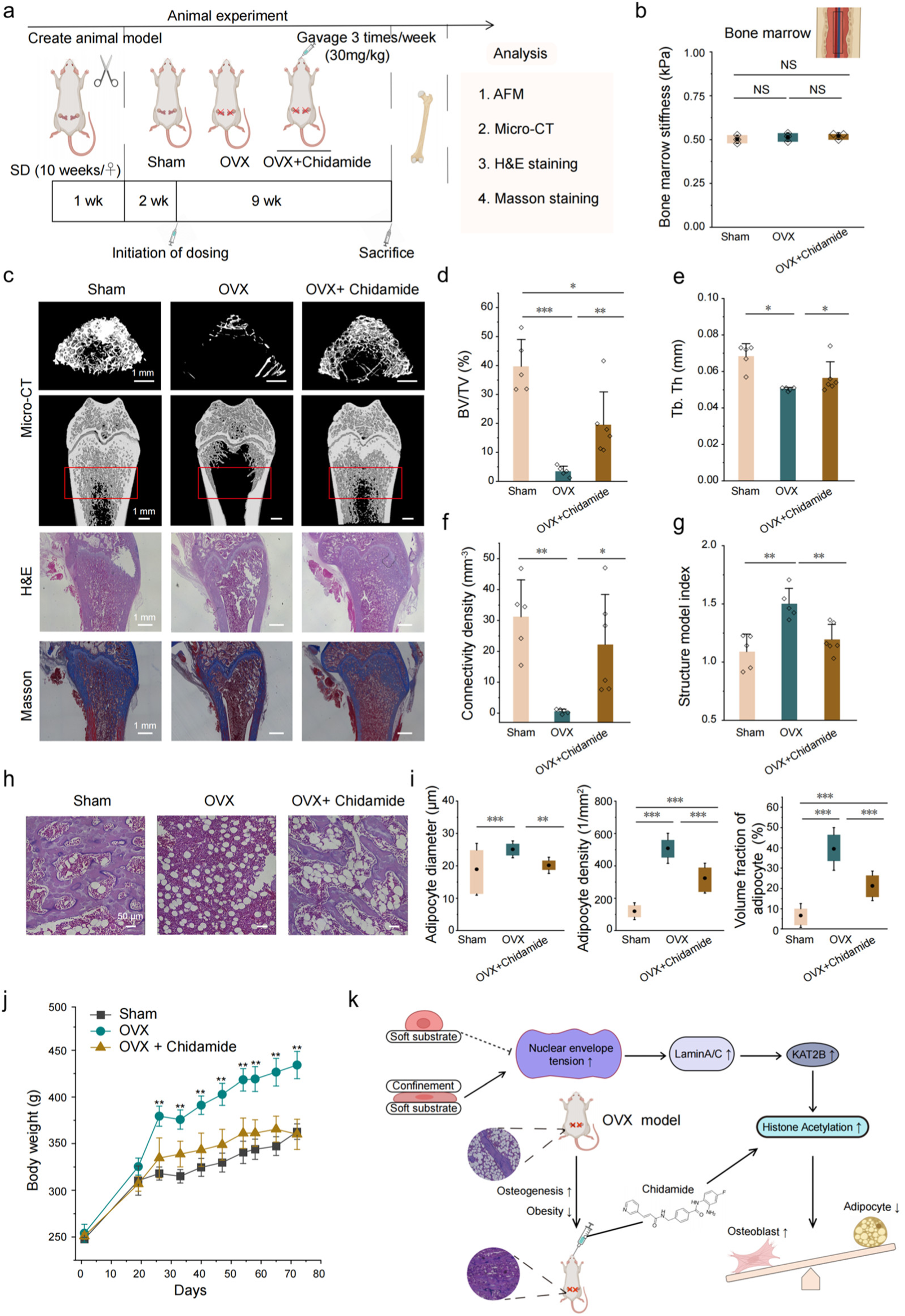
Pharmacological enhancement of histone acetylation effectively mitigates osteoporosis and obesity progression in rat models. (**A**) Schematic showing animal experiments: 10-week-old SD rats were adaptively fed for 1 week, OVX surgery to establish an osteoporosis model, Chidamide administration (30 mg/kg, 3 times a week) began 2 weeks after surgery, and was sampled after 9 weeks, bone marrow stiffness (AFM), bone microstructure (Micro-CT), and histology (HE/Masson) analysis. (**B**) Bone marrow stiffness of the sham operation group (Sham) (n=5), OVX model group (n=5) and OVX + Chidamide group (n=6). (**C**) Three-dimensional reconstruction of the femur Micro-CT, HE staining and Masson trichromatic staining of the Sham group, OVX model group and OVX + Chidamide group. (**D-G**) Bone volume fraction (BV/TV) (D), Trabecular bone thickness (Tb.Th) (E), Connection density (Conn.D) (F) and Structural Model Index (SMI) (G) of the Sham group (n=5), OVX model group (n=5) and OVX + Chidamide group (n=6). (**H**) HE staining of the adipocyte region of the Sham group, OVX model group and OVX + Chidamide group. (**I**) Average diameter of fat cells, adipocyte density and volume fraction of adipocytes of the Sham group, OVX model group and OVX + Chidamide group (4 biologically independent experiments in each group). (**J**) Weight temporal changes of the Sham group (n=5), OVX model group (n=5) and OVX + Chidamide group (n=6). (**K**) Schematic showing confinement-induced nuclear mechanotransduction mechanism. Data: B, C, E, F, G, H, K, mean± SD; B, C, G, H, K Tukey’s test; E, F Mann-Whitney test; ns, P > 0.05; *P ≤ 0.05; **P ≤ 0.01; ***P ≤ 0.001.

Notably, microcomputed tomography (micro-CT) evaluation, H&E and Masson’s trichrome staining demonstrated substantial bone loss in OVX rats, while Chidamide treatment group exhibited robustly enhanced bone repair (Fig. 5c). Quantitative 3D morphometry showed significant increases in bone volume fraction (BV/TV) (Fig. 5d) and femoral trabecular thickness (Tb.Th) (Fig. 5e) in Chidamide treatment group compared to OVX group (p<0.01). Elevated connectivity density (Fig. 5f) coupled with reduced structure model index (SMI) (Fig. 5g) further confirmed the drug’s capacity to maintain bone structural integrity and effectively reduce osteoporosis-related fracture risks.

On the other hand, femoral histomorphometry analyses using H&E staining of rat femur section demonstrated that the increases in the diameter, density, and volume fraction of adipocytes observed in OVX rats were effectively reduced after Chidamide treatment, corroborating Chidamide efficacy in preventing excessive adipogenesis induced by PMOP (Fig. 5h–i). Moreover, OVX model rats exhibited significant weight gain, whereas Sham and Chidamide-treated groups maintained relatively stable body weights (Fig. 5j), further indicating that pharmacological enhancement of histone acetylation can inhibited the PMOP-associated obesity.

Collectively, these data demonstrate that pharmacological enhancement of histone acetylation can effectively mitigates osteoporosis and obesity progression *in vivo*, highlighting the potential of nuclear mechanotransduction signaling as a novel therapeutic target for PMOP treatment (Fig. 5k).

## Discussion

### Spatial confinement induces BMSCs cultured on soft substrate to exhibit lineage commitment similar to those on unconfined stiff substrates

In fact, cells typically develop adaptive mechanisms to cope with the mechanical microenvironment *in vivo*, enabling self-protection and physiological functions [43]. As the most well-recognized biomechanical cue, ECM stiffness profoundly influences cellular behaviors such as spreading, migration, and differentiation [44]. This conventional stiffness-involved mechanotransduction mechanism cannot fully account for the osteogenic commitment of BMSCs within the soft bone marrow microenvironment *in vivo*, while our findings demonstrate that spatial confinement alone, without altering the mechanical properties of substrate, can induce BMSCs cultured on soft substrate to exhibit lineage commitment similar to those on unconfined stiff substrates, including adipogenesis suppression and osteogenesis promotion. This parallels our previous discovery that crowding-induced confinement makes quiescent round cells on soft adhesive substrates spread and migrate, mirroring behaviors typically observed on stiff substrates [29]. These results demonstrate that crowding-induced confinement can functionally override or compensate cell mechanosensing of ECM stiffness to some extent, highlighting that spatial confinement play an underappreciated yet pivotal role in regulating cellular behavior within native tissue microenvironments.

### Nuclear mechanosensing drives confinement-induced osteogenic differentiation via an epigenetically regulated pathway, independent of cell-substrate adhesion and YAP nuclear translocation

Previous studies have demonstrated that osteogenic differentiation of BMSCs is coordinately regulated by multiple signaling pathways, with ECM adhesion-dependent mechanotransduction playing a pivotal role. This classical paradigm posits that focal adhesion-mediated stress fiber contraction activates the RhoA/ROCK pathway and Yes-associated protein (YAP) nuclear translocation, thereby upregulating the expression of osteogenic master genes such as Runx2 and BMP2 [45, 46]. This mechanism heavily relies on cell-substrate adhesion, while our findings reveal that mechanical confinement can still promote Runx2 nuclear translocation and osteogenic differentiation even upon the inhibition of cell-substrate adhesion (Fig. 2a-c and Supplementary Fig. 3a-c). Our results establish nuclear mechanosensing—mediated through a Lamin A/C-dependent pathway that activates KAT2B and modulates histone acetylation—as a primary and adhesion-independent driver of confinement-induced osteogenesis (Fig. 6). This discovery aligns with prior studies on nuclear tension-mediated gene regulation [47], highlighting the nucleus as an autonomous mechanical sensor.

**Fig. 6:**
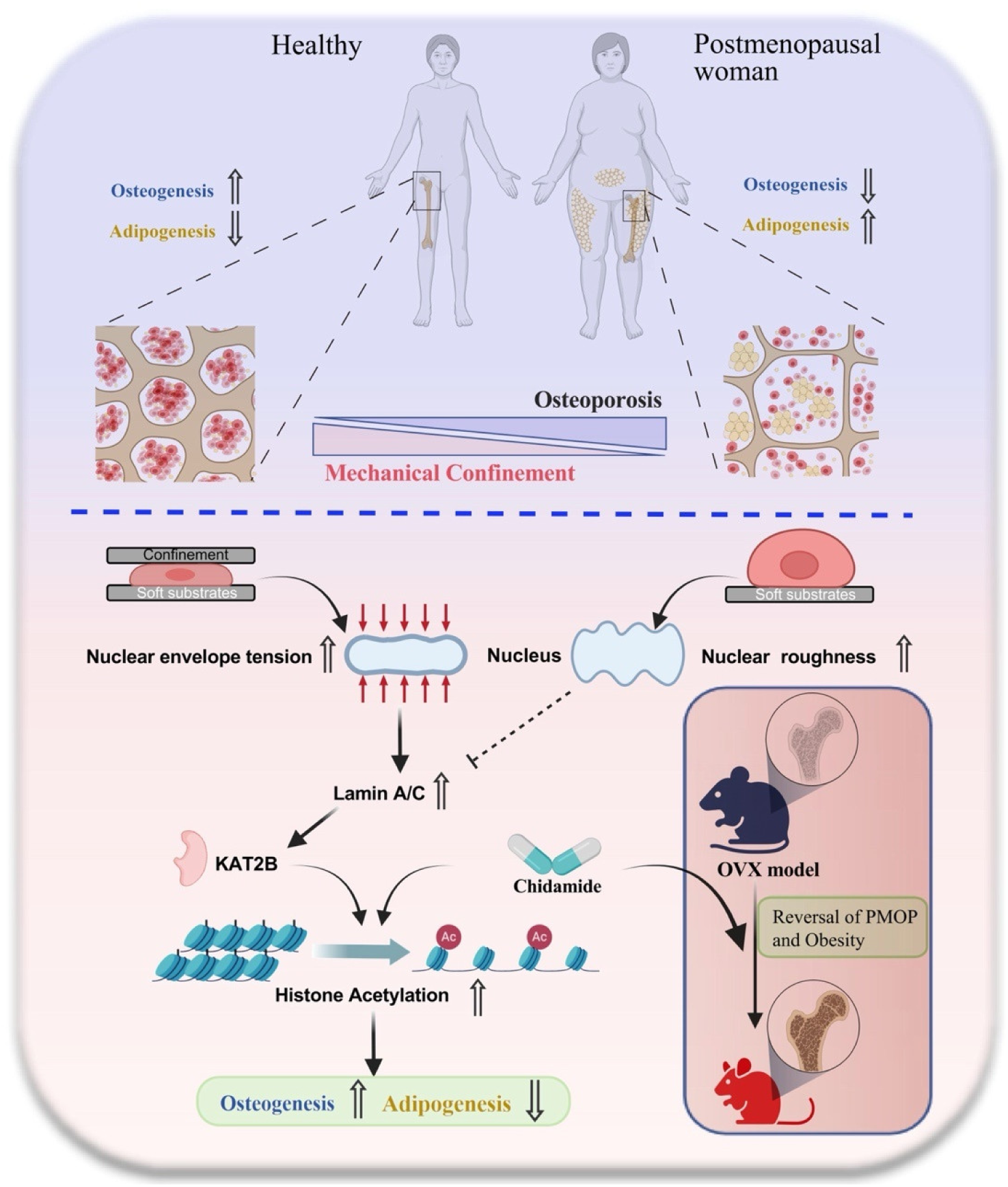
Schematic showing the crowding-induced nuclear mechanical remodeling inhibiting postmenopausal osteoporosis and obesity via KAT2B-mediated histone acetylation.

Indeed, existing studies also indicate that nuclear envelope stretching can induce conformational changes in nuclear pore complexes (NPCs), directly promoting YAP nuclear transport without cell-substrate adhesion [29, 34]. To further validate the independence of our mechanism from YAP signaling, we treated BMSCs with verteporfin (a YAP-TEAD interaction inhibitor). We found that even under complete YAP nuclear localization blockade (Supplementary Fig. 5a-b), spatial confinement maintained increased histone acetylation (Supplementary Fig. 5b), high Runx2 nucleus/cytoplasm ratio and osteogenic differentiation capacity (Supplementary Fig. 5c-d). Thus, we propose that confinement-driven osteogenesis depends on nuclear mechanosensing but is mainly regulated by epigenetic histone acetylation, independent of YAP nuclear translocation. Moreover, a recent study has reported that mechanical confinement promotes the transition of melanoma cells from proliferative to drug-resistant invasive types through chromatin remodeling [48], indicating such confinement-induced epigenetic modifications may have broader implications and influence diverse cellular physiological behaviors.

### Nuclear mechanosensing governs both acute and persistent cellular responses

Emerging evidence highlights the nucleus as a mechanosensor capable of transducing extracellular mechanical cues to regulate cellular behaviors. For instance, nuclear deformation can trigger rapid migratory transitions in quiescent cells [29, 49–51], or promote mature focal adhesion formation to enhance cell-substrate adhesion [29], demonstrating the pivotal role of nuclear mechanotransduction in the regulation of short-term cellular behaviors. However, beyond these short-term responses (e.g., adhesion and migration), our work reveals its capacity to orchestrate long-term cell fate decisions. We demonstrate that confinement-induced nuclear deformation leads to NE tension elevation and Lamin A/C-dependent activation of KAT2B, triggering persistent histone acetylation upregulation that commit cells to osteogenic differentiation on long-term time scales (Fig. 6). This perspective is further corroborated by emerging evidence that nuclear mechanotransduction can directly influence enduring biological processes, including cellular reprogramming and lineage specification [45, 52–55], underscoring the broad relevance of nuclear deformation as a regulator of cell identity and function. Therefore, nuclear mechanosensing regulate not only the short-term cellular behaviors, but also exhibit “hysteresis”, enabling cells to retain mechanical memory over extended durations.

### Nuclear mechanotransduction as a therapeutic target: implications for bone repair and beyond

Our findings demonstrate that enhancing histone acetylation through Chidamide and reversing nuclear mechanical signaling transduction can significantly improve bone regeneration and inhibit obesity in PMOP model rats. Given that the nucleus serves as the cell’s primary mechanosensory hub—integrating biochemical, physical, and mechanical inputs to orchestrate diverse cellular functions—its mechanical regulation extends far beyond osteogenesis. Indeed, dysregulated nuclear mechanotransduction underpins a spectrum of pathologies, from laminopathies to metastatic cancer [55]. For instance, Hutchinson-Gilford progeria syndrome (HGPS), driven by Lamin A/C mutations, disrupts nuclear integrity and accelerates chromatin aging via aberrant H3K27me3 redistribution [56]. Similarly, Lamin A/C mutations cause dilated cardiomyopathy (DCM) with lethal arrhythmias, highlighting the cardiac dependence on nuclear mechanical homeostasis [57]. Beyond genetic disorders, ECM stiffness promotes tumor progression by increasing Lamin A/C-mediated nuclear tension, thereby facilitating YAP-dependent oncogenic transcription [58]. Therapeutically, targeting nuclear mechanics has shown clinical promise: Lonafarnib (Zokinvy), an inhibitor of farnesylation of the Lamin A precursor, became the first FDA-approved therapy to prolong survival in HGPS patients [59]. Thus, the confinement-mediated nuclear mechanotransduction mechanism we found here may be important for a wide range of physiological and pathological processes, including osteoporosis and obesity, and is a promising target for clinical intervention. These insights position nuclear mechanosensing as a pivotal regulator of physiology and disease, offering a compelling avenue for mechano-epigenetic interventions in regenerative medicine and beyond.

### Chidamide: a new candidate for holistic treatment of PMOP and concomitant obesity

Current clinical strategies for PMOP usually focus solely on either inhibiting bone resorption or promoting bone formation—each constrained by significant limitations. For example, anti-resorptive agents such as zoledronic acid can effectively reduce the fracture risk of osteoporotic patients [60], but their use may be associated with side effects including gastrointestinal intolerance, nephrotoxicity, and even osteonecrosis [61, 62]. As for anabolic agents like teriparatide, although effective in alleviating osteoporosis, their application is highly limited due to the risk of osteosarcoma (with treatment duration strictly restricted in 2 years) and the requirement for daily injections, which adversely affects patient compliance [13]. Critically, none of the existing clinical strategies address the frequent PMOP-associated obesity.

In stark contrast, we demonstrate that Chidamide—an orally available deacetylase inhibitor approved by FDA—simultaneously promotes osteogenesis and suppresses adipogenesis by modulating nuclear mechanotransduction (Fig. 6). This dual curative effects positions Chidamide as a novel candidate for holistic treatment of PMOP and concomitant obesity, potentially redefining therapeutic standards for metabolic bone diseases. Furthermore, its oral administration offers superior convenience compared to injectable alternatives. Further controlled clinical trials in postmenopausal populations are needed to further validate its efficacy and safety profile.

## Methods

### The working concentrations for all antibody reagents utilized are provided in Table 1

#### Cell confinement device fabrication

The cell confinement device lid comprises three components: (1) a thin polydimethylsiloxane (PDMS) sheet featuring an array of micropillars of defined height, (2) an 18-mm diameter glass cylinder, and (3) a complementary lid structure designed to hold the glass cylinder. To fabricate this PDMS sheet, a silicon wafer master mold was used, patterned with a regular array of holes (diameter: 0.44 mm, pitch: 1 mm) via standard photolithography. Polydimethylsiloxane (PDMS) prepolymer (Sylgard 184, Dow Corning) was prepared by thoroughly mixing the base (Part A) and curing agent (Part B) at an 8:1 (w/w) ratio. The PDMS mixture was degassed, then poured onto the silicon master mold and cured in an oven at 65°C for 12 hours. After curing, the solidified PDMS sheet was carefully peeled from the mold and cut to the appropriate dimensions (this sheet contains an array of micropillars of defined height). The 18-mm diameter glass cylinder and the PDMS micropillar sheet were subjected to oxygen plasma treatment (PDC-002, Harrick, Stockton, CA) for 60 seconds. Immediately after plasma activation, the PDMS micropillar sheet was irreversibly bonded to the flat surface of the glass cylinder, ensuring precise alignment. The assembled lid structure was then fitted into its complementary holder (a component designed to hold the glass cylinder), completing the fabrication of the cell confinement device lid.

Polyacrylamide hydrogels with tunable stiffness were prepared on glass-bottom confocal dishes (NEST or Biosharp) following established protocols, with modifications as detailed below. Glass-bottom confocal dishes were treated sequentially with 3-Aminopropyl trimethoxysilane in ddH₂O for 15 minutes and 0.05% (w/v) glutaraldehyde in phosphate-buffered saline (PBS) for 30 minutes. Dishes were rinsed thoroughly with distilled water and ethanol after each step and dried under nitrogen gas. Glass coverslips used for gel casting were cleaned and silanized by immersion in Trimethylchlorosilane in ethanol for 15 minutes, followed by extensive rinsing with ddH₂O, and drying under nitrogen. PA gel precursor solutions of desired stiffness were prepared by mixing appropriate volumes of the following stock solutions in microcentrifuge tubes: 40% (w/v) acrylamide, 2% (w/v) N,N’-methylenebisacrylamide (bis-acrylamide), distilled water, 1% (w/v) ammonium persulfate (APS), and 0.1% (v/v) N,N,N’,N’-Tetramethylethylenediamine (TEMED). Final concentrations of acrylamide and bis-acrylamide determined the gel stiffness. 90 µL of the freshly mixed precursor solution was pipetted onto the silanized surface of a confocal dish. A Trimethylchlorosilane-silanized coverslip was carefully placed on top of the droplet, ensuring no air bubbles were trapped between the coverslip and the solution. Polymerization was allowed to proceed undisturbed at room temperature for 40 minutes.

After polymerization, 1 mL of distilled water was gently added to the dish surrounding the gel. The dish was incubated for 15 minutes at room temperature to hydrate the gel. The top coverslip was then carefully removed using forceps. The hydration water was aspirated and replaced with 1 mL of fresh distilled water. Hydrated PA gels were stored at 4°C until further use (typically within 1 week). Gel surfaces were activated for protein conjugation using the heterobifunctional crosslinker Sulfo-SANPAH. 110 µL of 0.25 mg/mL Sulfo-SANPAH solution in distilled water was applied to the gel surface. The gel was exposed to UV light (365 nm) for 15 minutes to photoactivate the Sulfo-SANPAH. The gel was washed three times with 50 mM HEPES buffer (pH 8.5). Fibronectin (FN) solution (10 µg/mL in PBS) was applied to the activated gel surface and incubated overnight at 4°C. Prior to cell seeding, the FN solution was aspirated, and the functionalized gel surface was rinsed thoroughly with PBS.

Prior to each experiment, the PDMS micropillar surface of the confinement device lid was incubated with 0.1% (w/v) Pluronic F-127 (Thermo Fisher Scientific) in 10 mM HEPES buffer (pH 7.4) for 1 hour at room temperature to prevent non-specific cell adhesion. The F-127 solution was aspirated, and the device lid was equilibrated by incubation in complete cell culture medium for at least 30 minutes at 37°C. Cells were seeded onto the FN-functionalized PA gels within confocal dishes and allowed to adhere and spread under standard culture conditions for the desired duration. The device lid was slowly and vertically lowered onto the center of the cell-seeded PA gel, applying gentle, consistent pressure to ensure full contact between the micropillars and the gel surface, thereby deforming the underlying cells. Confinement was maintained throughout the observation period.

#### Cell culture and induced differentiation

Human adult mesenchymal stem cells from bone marrow and corresponding induction media were purchased from Cyagen Biosciences. Cells were routinely maintained in low-glucose Dulbecco’s Modified Eagle Medium (DMEM) supplemented with 10% (v/v) fetal bovine serum (FBS), 1% (v/v) L-glutamine, 1% (v/v) penicillin-streptomycin. Cells were cultured at 37°C in a humidified atmosphere containing 5% CO₂, and were subcultured when they reached 80–90% confluence. All experiments utilized cells at or before passage 5 (P5). Preparation of induction medium: Osteogenic Induction Medium and Adipogenic Induction Medium were mixed at a 1:1 (v/v) ratio to prepare Osteo-Adipogenic Mixed Induction Medium (OAMIM). BMSCs were seeded onto fibronectin-functionalized polyacrylamide (PA) gel substrates prepared in glass-bottom dishes (as detailed in the “Preparation of Polyacrylamide Hydrogel Substrates” section). After culturing in complete growth medium for at least 24 hours to ensure full cell adhesion, the medium was aspirated, and pre-mixed OAMIM was then aseptically added to the dishes. Cells were incubated under standard culture conditions (37°C, 5% CO₂) for 6 hours before mechanical stimulation. Cells were seeded at a density of 4 × 10³ cells per dish (35-mm-diameter culture dishes) and were allowed to adhere overnight. For drug intervention experiments, the concentrations and batch numbers used were listed in the Key Resource Table. Drugs were administered via a confinement device for 6-12 hours, and the culture medium were replaced every two days. After a period of culture, cells were fixed for staining and observation.

#### Staining and imaging analysis

Without specific indication, Bright-field staining of ALP of BMSCs and drug treatment experiments are generally collected after 7 days culture, while immunofluorescence staining is collected after 3 days culture. Cells were fixed with 4% (w/v) paraformaldehyde (PFA) in phosphate-buffered saline (PBS) for 15 min at room temperature (approximately 25°C), followed by 3 washes with PBS (5 min each). Fixed samples were permeabilized with 0.25% (v/v) Triton X-100 in PBS for 15 min, and non-specific binding sites was blocked with 5% (v/v) normal goat serum in PBS for 1 h at room temperature. Samples were then incubated with primary antibodies for 2 h. After 3 washes with PBS (5 min each), samples were incubated with fluorophore-conjugated secondary antibodies (dilution in blocking solution) for 1 h in the dark. Following secondary antibody incubation, samples were washed 3times with PBS (5 min each) and stained with 4’,6-diamidino-2-phenylindole (DAPI) for 30 min at room temperature in the dark. Final washes were performed with PBS (3 washes, 5 min each). Images were acquired using a Leica SP8 confocal laser-scanning microscope (Leica Microsystems, Germany) equipped with a 40× water-immersion objective (numerical aperture [NA]: 1.1). Z-stacks were collected at 1 μm intervals through the entire cell volume. Laser power, gain, and offset were optimized for each channel to avoid saturation and maintained constant across compared samples. Z-stack reconstruction: Maximum-intensity projections (MIPs) of individual cells were generated using the Z-Project function in ImageJ (National Institutes of Health, USA) or Leica Application Suite X (LAS X). The nuclear/cytosolic fluorescence intensity ratios of YAP and Runx2 were quantified by measuring the mean fluorescence intensity in the nucleus (defined by DAPI staining) and cytoplasm (defined as the cellular area excluding the nucleus) using ImageJ.

#### Bulk RNA sequence

Total RNA was isolated from bone marrow mesenchymal stem cells (BMSCs) using TRIzol™ reagent (Thermo Fisher, Cat# 15596018) according to the manufacturer’s protocol. RNA quantity and purity were assessed using a Qubit™ 3.0 Fluorometer (Thermo Fisher, Cat# Q33216) and an Agilent 5300 Fragment Analyzer (Agilent Technologies, Cat# M5311AA). High-quality RNA samples (RNA Integrity Number, RIN > 7.0) were used for library construction. Sequencing libraries were prepared from pooled human RNA samples. The resulting cDNA libraries were sequenced on an Illumina NovaSeq™ X Plus platform using a paired-end (2×150 bp) RNA-seq approach. Differential gene expression analysis was performed using DESeq2 for group comparisons and edgeR for pairwise sample comparisons in R software. Genes with a false discovery rate (FDR) < 0.05 and an absolute fold change ≥ 2 were defined as differentially expressed genes (DEGs). DEGs were analyzed using Gene Ontology (GO) functional enrichment analysis (biological process, cellular component, and molecular function categories) and Kyoto Encyclopedia of Genes and Genomes (KEGG) pathway enrichment analysis, implemented in clusterProfiler with the latest GO database (release 2023-05) and KEGG database (release 106). Gene Set Enrichment Analysis (GSEA) was also performed to identify enriched gene sets. The raw sequence data reported in this paper have been deposited in the Genome Sequence Archive (Genomics, Proteomics & Bioinformatics 2021) in National Genomics Data Center (Nucleic Acids Res 2025), China National Center for Bioinformation/Beijing Institute of Genomics, Chinese Academy of Sciences (GSA-Human: HRA012498) that are publicly accessible at https://ngdc.cncb.ac.cn/gsa-human [63, 64].

#### Transfections

Cells were seeded into 6-well confocal microplates containing soft substrates. Transfection was performed when cell confluence reached approximately 60%. The transfection reagent (Genepharma, Cat# G04026) was used, including Buffer and Plus reagent. For each transfection reaction, 28 μL of Buffer was added to a sterile, RNase-free microcentrifuge tube. Then, 50 pmol of siRNA (2.5 μL of a 20 μM stock solution) was added, and the mixture was gently pipetted to form the siRNA pre-mix. Next, 10 μL of Plus reagent was added to the siRNA pre-mix, followed by immediate and thorough mixing via pipetting up and down several dozen times to form the siRNA/Plus complex. The prepared siRNA/Plus complex was added to the cells, and the microplate was gently shaken 10-15 times in both front-back and left-right directions to ensure uniform distribution. Cells were then returned to the incubator for continued culture. A mechanical confinement device was applied 12 hours post-transfection. Protein level changes were detected 72 hours post-transfection. For assessment of osteogenic differentiation, alkaline phosphatase (ALP) activity was measured at 7 days post-transfection. All cells received an additional siRNA supplement on day 3 after transfection.

#### Atomic force experiments and quantification

AFM experiments were conducted using an AFM (BioScope Resolve 8, Bruker, Germany) mounted on top of a Leica DMi8 fluorescence microscope. Polystyrene microspheres with a diameter of 20 μm were attached to the end of a blunt-tipped NP-O10 cantilever using a non-fluorescent adhesive. The laser spot was aligned to the free end of the cantilever using the system’s laser alignment module, with fine adjustments made via the beam positioning knob. Cantilever deflection sensitivity was calibrated via thermal tuning according to the manufacturer’s protocol, enabling sufficient mechanical stimulation to selectively deform the measurement via contact mode. For polyacrylamide (PA) hydrogel stiffness measurements, the AFM probe (0.12 N/m cantilever with a 10 μm-radius polystyrene microsphere, equivalent to 20 μm diameter) generated force-distance (F-D) curves using consistent parameters: loading peak force (5 nN), retraction speed (10 μm/s), and measurement range (5 μm). For each sample, at least 5 fields of view were randomly selected, with at least 9 F-D curves collected per field of view. The modulus was calculated from F-D curves using Bruker NanoScope Analysis software (v3.0), with stiffness fitted via the DMT model. The fitting force range was restricted to the 5%–50% interval after the contact point, and Poisson’s ratio was set to 0.5.

For rat bone marrow stiffness measurements, the rat femur was rapidly dissected after anesthesia. Each femur shaft was split longitudinally along its long axis to expose the bone marrow (including the central vein), and samples were immediately immersed in pre-chilled PBS to maintain viability. AFM indentation measurements were performed at multiple distinct locations on the exposed bone marrow surface. Sampling parameters were identical to those used for PA hydrogel measurements (see above). Prior to measurements, AFM probes were pre-treated by soaking in 0.1% (w/v) Pluronic F-127 overnight to minimize non-specific adhesion.

#### Quantitative real-time polymerase chain reaction

Total RNA was extracted from mesenchymal stem cells using an RNA isolation kit (Cat# AC0205-A, SparkJade), and its concentration was determined using NanoDrop One (Thermo Fisher). PrimeScript RT premix (RR036A, Takara Bio) was used, followed by SYBR qPCR premix (Cat# 22204, Tolo Biotech) for qRT-PCR. PCR reactions were performed on the LightCycler 96 instrument (Roche LifeScience), and relative mRNA expression was calculated using the 2^(-ΔΔCt) method. The gene-specific primers used are listed in the table.

#### Western blotting

Western blot analysis was performed following standard protocols with modifications. Briefly, cells were lysed in 5× protein loading buffer (containing DTT). After centrifugation and heat denaturation, the lysate was loaded onto a 12% polyacrylamide gel (SDS-PAGE Gel Quick Preparation Kit, Beyotime) for electrophoresis-based protein separation and transferred to a polyvinylidene fluoride (PVDF) membrane (Millipore); Prepare a 5% milk blocking solution and incubate the membrane for 90 minutes. After blocking, wash the membrane with TBST and incubate overnight at 4°C with the primary antibody (Lamin A/C antibody diluted 1:2000, H3 antibody diluted 1:1000) at 4°C overnight. Then, the membrane was washed and incubated with horseradish peroxidase (HRP)-conjugated secondary antibody (diluted 1:30000) at room temperature for 2 hours. The ECL Chemiluminescence Substrate Kit (CYTOCH) was used for color development, and the bands were visualized using the Amersham ImageQuant 800 system (Cytiva). Band intensity was analyzed using ImageJ software.

#### Scanning electron microscopy

Pre-prepared soft and stiff polyacrylamide (PA) hydrogels were immersed in deionized water and equilibrated for 15 min at room temperature under sealed conditions. To induce brittle fracture, samples were rapidly frozen in liquid nitrogen (-196°C) for 10 min. Fractured hydrogel fragments were immediately transferred to a freeze-dryer (Scientz) and vacuum-dried at -50°C under a pressure of <10 Pa for 48 h. Conductive carbon adhesive tape was affixed to aluminum SEM stubs. Lyophilized fragments were handled with anti-static tweezers and firmly mounted onto adhesive surfaces. Prior to SEM observation, samples were sputter-coated with a 5-nm-thick gold layer using an ion sputter coater (Leica EM ACE600, Leica Microsystems) under an argon atmosphere at a current of 30 mA for 100 s. Finally, SEM images were acquired using a SEN5000 scanning electron microscope (Hitachi High-Tech).

#### Animal experiments

For the OVX model, 10-week-old female Sprague-Dawley (SD) rats were purchased from Charles River Laboratories International, Inc. (NYSE:CRL). The rats underwent either bilateral ovariectomy to simulate postmenopausal osteoporosis or a sham surgery. One week after arrival, the surgical procedures were performed. Prior to surgery, the rats were fasted for 12 hours with free access to water. Anesthesia was induced by intraperitoneal injection of a freshly prepared anesthetic solution (10 ml/kg), with a new batch prepared for every two rats. The abdominal area was shaved. The surgical site was disinfected with an iodine solution. A small incision (approximately 1–2 cm in length) was made along the midline of the abdomen and the base of the hind legs. The abdominal cavity was opened, and one side was clamped with a hemostat. The uterus was identified, and the ovaries, which are cauliflower-shaped and approximately the size of a mung bean, were located on each side via the ‘Y’ bifurcation of the uterus. The uterus was ligated at a lower position, followed by ligation of the junction between the ovary and the uterus along with the surrounding fat tissue using a 5-0 suture. Both ovaries were completely excised. Minor bleeding, if present, was controlled with cotton compression. For the sham surgery group, the identical surgical procedure was performed, including the location of the ovaries, but without ligation or removal; only a small amount of adjacent fat tissue was excised. All procedures were performed on a heating pad. After suturing the muscle layer, the wound was cleaned with iodine solution to remove blood stains and treated with gentamicin. The epidermis was then sutured, and the wound was disinfected again with iodine solution. All animal care and experimental procedures were conducted in strict accordance with the guidelines in the Guide for the Care and Use of Laboratory Animals and were approved by the Animal Care and Use Committee of the Institute of Health and Medicine, Hefei Comprehensive National Science Center (Approval ID: IHM-CLAM-AP-2025-003).

#### Bone micro-CT analysis

Following fixation in 4% paraformaldehyde for 48 hours, the femora were scanned using a micro-CT system (Imaging 100, Raycision Medical Technology Co., Ltd) at 45 kVp, 250 μA, and with an isotropic resolution of 10 μm. The resulting images were segmented, and the regions of interest (ROIs) were analyzed using the device’s proprietary software. For the assessment of trabecular bone microstructure, a 1.5 mm-long ROI was selected, commencing 0.5 mm proximal from the distal growth plate to avoid the primary spongiosa. Three-dimensional reconstructed models were generated to visualize the trabecular architecture.

#### Bone histology

Human bone specimens were sectioned into approximately 5 mm-thick slices and decalcified by immersion in 10% (w/v) EDTA (pH 7.4) at room temperature. The EDTA solution was replaced regularly (2–3 times per week). The endpoint of decalcification was determined by a needle penetration test, whereby a fine needle was gently pressed against the densest cortical area of the sample. Decalcification was considered complete when the needle could penetrate the tissue without significant resistance. The duration of decalcification depended on sample size and density, typically requiring 1 to 2 weeks. Human bone specimens were kindly provided by L.W. (The First Affiliated Hospital of the University of Science and Technology of China). All procedures involving human samples were approved by the Ethics Committee of The First Affiliated Hospital of the University of Science and Technology of China (No. 2025KY429).

Following euthanasia, rat femora were promptly dissected. All attached soft tissues (e.g., muscle and fascia) were meticulously removed to facilitate reagent penetration. A longitudinal incision was made along the long axis of the bone shaft to expose the internal architecture, including the cortical bone, medullary cavity, and growth plate. The samples were first fixed by immersion in a sufficient volume of neutral buffered formalin (at least 10 times the tissue volume) for 24–72 hours. Subsequently, the fixed tissues were decalcified by complete immersion in a decalcification solution consisting of 10% formic acid and 10% formalin (volume at least 20–50 times that of the tissue). Beginning on day 3-5 of decalcification, the endpoint was assessed daily using a needle penetration test, preferentially applied to the cut surface or areas of thinner cortical bone. Upon reaching the endpoint, tissues were immediately removed from the decalcification solution and rinsed thoroughly under running water. The decalcified bones were then processed through routine histological procedures, including dehydration through a graded ethanol series, clearing in xylene, and paraffin embedding. Sections (3–5 μm thick) were prepared and stained using either a Hematoxylin and Eosin (H&E) staining kit (Beyotime, #C0105S) or a Masson’s trichrome staining kit (Beyotime, #C0189S). Stained sections were imaged using a Thunder imaging system.

#### Human samples

Bone marrow and femur were obtained from The First Affiliated Hospital of USTC, Division of Life Sciences and Medicine, University of Science and Technology of China. All human samples used in this study were approved by the Medical Ethics Committees of The First Affiliated Hospital of USTC, Division of Life Sciences and Medicine, University of Science and Technology of China. (2025 KY no.429).

#### Statistical analysis

All data were analyzed using Origin or Leica Application Suite X (LAS X). Results are presented as means ± SEM. Unless otherwise specified, an unpaired two-tailed t test was used for comparisons between two groups. Statistical significance was determined at a P value threshold of < 0.05, In all figures, measurements are reported as Mean ± SD. The data analyzed were pooled from many cells/fields of view over several fields of view across n ≥ 3 experiments. The number of data points for each experiment and significance levels are noted in the figure text.

## Material availability

This study did not generate any materials and unique reagents.

## Acknowledgments

This work was supported by the National Natural Science Foundation of China (Grants No. 12025207 and No. 12532015), the Fundamental Research Funds for the Central Universities, and USTC Research Funds of the Double First-Class Initiative (YD2090002024, YD2090002502). This work was partially carried out at the University of Science and Technology of China Center for Micro and Nanoscale Research and Fabrication.

## Author Contributions

H.J., H.Y. conceived and conducted the project; H.J., H.Y. and M.X. designed the experiments; L.W. collected human samples and clinical information; M.X. performed main experiments; W.W and H.Y. participated in parts of experiments; M.X and H.Y. summarized the data; H.J., H.Y. and M.X. wrote the manuscript; H.J. supervised the project.

